# Shifts in consumer assemblages are linked to nutrient availability and ecosystem metabolism

**DOI:** 10.64898/2026.04.23.720454

**Authors:** Isabella G. Sadler, Adrienne L. Stanley, Charlotte F. Narr

## Abstract

Nutrient availability, ecosystem productivity, and consumer assemblages are intricately linked through complex interactions and feedbacks. Nutrients influence the diversity and functional roles of consumers via shifts in resource quality and quantity, and consumers can alter ecosystem production and nutrient availability. However, our understanding of how characteristics of consumers respond to and influence concomitant shifts in nutrient availability and production is limited. We quantified the response of well-studied consumer assemblages (benthic invertebrates and zooplankton) to realistic nutrient loads that altered gross primary production (GPP) and ecosystem respiration (ER). We fertilized 14 outdoor experimental ponds for 2 months and monitored total water column carbon (TC), nitrogen (TN), and phosphorus (TP), GPP, ER, and net ecosystem production (NEP) weekly. Then, we evaluated how fertilization and the variation in nutrients and metabolism caused by fertilization were related to shifts in consumer assemblages. Fertilization increased water column TN and TP and reduced TC:TP ratios, TN:TP ratios, and rates of GPP and ER. However, consumer assemblages were more tightly linked to variation in nutrient availability and production across ponds than to fertilization. Greater declines in benthic diversity occurred in ponds with higher average TN:TP ratios during the experiment. Consistent with predicted effects of cladocerans on nutrient availability, shifts in cladoceran abundances were positively associated with average water column TN:TP ratios during the experiment. Finally, elevated GPP and ER were associated with greater increases in the abundance of benthic invertebrate predators, suggesting the possibility of top-down control. Our study highlights the critical role of consumer-mediated processes in the interaction between nutrient availability and production.

**Manuscript Highlights:**

- Fertilization reduced pond gross primary production and ecosystem respiration rates.
- Invertebrate predator abundance was inversely related to gross primary production.
- Shifts in consumer assemblages were tightly linked to nutrients and production.

## Introduction

Nutrient availability influences many ecosystem processes, often limiting primary production (Hecky and Kilham 1988, Elser et al. 2007). Increases in nutrients like nitrogen (N) and phosphorus (P) can stimulate rates of ecosystem metabolism, such as gross primary production (GPP), ecosystem respiration (ER), and net ecosystem production (NEP) (Hanson et al. 2003, Staehr et al. 2010). Changes in both nutrient availability and ecosystem metabolism alter biodiversity within ecosystems (Alimov 2001, Rader & Richardson, 1992,Wang et al.2021). However, the direction of these shifts is difficult to predict (Elser et al. 2007, Halpern et al. 2008). Quantifying the effects of nutrient availability on ecosystem metabolism and biodiversity is increasingly important as anthropogenic activities continue to increase nutrient loading in freshwaters (Vitousek et al. 1997, Peñuelas et al. 2012) and concerns about global carbon cycling grow (del Giorgio and Peters 1994, Balmer and Downing 2011).

Delineating the link between nutrient availability and production requires a better understanding of how their interaction is influenced by consumer assemblages (Welti et al. 2017). Nutrient availability and trophic interactions influence production (Schindler and Fee 1974, Carpenter et al. 1988), but consumer characteristics also feedback to influence nutrient availability (Elser and Urabe 1999). Experimental manipulation of nutrients and predators is a classic study design to understand these interactions (Brett and Goldman 1997). However, despite mounting evidence that animals can have large effects on both nutrient availability (Atkinson et al. 2017) and production (Rizzuto et al. 2024), characteristics of consumer assemblages are rarely incorporated into studies exploring the link between nutrients and ecosystem production. As a result, the role of consumer assemblage characteristics in mediating this link is poorly articulated, and predictions regarding the effects of nutrient inputs on production remain poorly parameterized. This lack of empirical data limits our understanding of which forces dominate these links in ecosystems.

Difficulty clarifying the role of specific consumers in this link is, in part, due to the challenge of classifying consumers in ways that accurately describe their trophic position and role within nutrient cycles, especially at the ecosystem scale. While the effects of nutrients and predators are often compared in a fully crossed design (Brett and Goldman 1997), the precise trophic position of many predators can be difficult to quantify, complicating efforts to quantify trophic cascades (Matich et al. 2011). Likewise, a growing body of literature demonstrates that consumer characteristics dictate both their response to and influence on basal nutrient availability, but studies that link consumer characteristics to nutrient availability at the ecosystem scale remain rare. One classification framework that could help overcome these issues is that of functional feeding groups (Cummins et al. 2005). This system uses morphological and behavioral characteristics that are functionally linked to benthic invertebrate modes of acquiring food, and, as a result, can be used to more rigorously classify entire freshwater invertebrate communities according to their trophic position and role within nutrient cycles (Cummins et al.2005).

Here, we work to understand how consumer traits mediate the relationship between nutrient availability and production by applying a classic experimental design (a fertilization experiment) to a well-studied system (invertebrate assemblages in experimental ponds) for which rigorous tools are available to evaluate nutrient availability and production, and the trophic positions of consumers can be characterized into functional feeding groups. We investigated the relationship between nutrient enrichment and ecosystem metabolism (GPP, R, NEP) by fertilizing 7 of 14 experimental ponds.

In addition to this direct assessment of fertilization effects, we also assessed how the variation in nutrient availability and ecosystem metabolism produced by fertilization was linked to shifts in benthic invertebrate diversity, the abundance of invertebrates in each functional feeding group, the quality and quantity of their diets (coarse particulate organic matter, CPOM, and fine particulate organic matter, FPOM), and the abundance of zooplankton within ponds. Within zooplankton, we focus on cladocerans and copepods, two taxonomic groups whose body size and nutritional requirements are often found to alter both their response to and effects on nutrient availability. We tested if the relationships we observed between each group of invertebrates and nutrient availability were triggered by our fertilization experiment through bottom-up effects, or if these relationships reflected more complex relationships that involved animal-mediated processes (e.g. consumer-driven nutrient cycling or trophic cascades). Given the morphological specificity of benthic invertebrate mouthparts, we anticipated that large populations of these invertebrate predators would produce more readily detectable trophic cascades in ecosystems where they were top predators. To assess this possibility, we compared how well basal resources explained shifts in consumer abundances using our entire dataset, which includes three ponds with large pre-existing populations of bluegill and green sunfish, and using the subset excluding these ponds.

We expected fertilization to enhance the quality and quantity of FPOM and stimulate gross primary production and that this stimulation of basal resources would increase benthic invertebrate diversity and the abundance of zooplankton and other filtering collectors that specialize on FPOM. We also anticipated that the relatively low N:P ratios of our fertilization treatment would stimulate the abundance of cladocera. Finally, we anticipated that shifts in the abundance of predacious benthic invertebrates and fish might alter production. We also suspected that large populations of fish with general morphological feeding structures (e.g., bluegill *Lepomis macrochirus* and green sunfish *Lepomis cyanellus*) will diminish our ability to detect relationships between basal resources and invertebrate assemblages. Overall, our work highlights how fertilization experiments in ponds not only allow for rigorous hypotheses testing in realistic conditions, but also provide critical information regarding an understudied, but important ecosystem within the global carbon cycle (Downing et al. 2006). Because small man made ponds are frequently fertilized to enhance the production of fish (Illinois Department of Natural Resources 2017), assessing how fertilization influences the net storage or loss of carbon via shifts in ecosystem metabolism could also help shape efforts to improve carbon storage.

## Methods

### Study Site

Our experiment was conducted in 14 approximately 0.04 hectare experimental ponds at a large outdoor research pond facility at Touch of Nature Environmental Center in Carbondale, IL. The study was conducted from May to October 2021. Existing amphibian, reptile, and/or fish communities were present in the ponds. Planktivorous fish species included *Lepomis macrochirus* (Bluegill) and *Lepomis cyanellus* (Green Sunfish). No piscivorous species were found in the ponds. Herpetofauna included *Chrysemys picta* (Painted Turtle), *Trachemys scripta elegans* (Red-eared Slider), *Chelydra serpentina* (Snapping Turtle), *Nerodia sipedon* (Common Watersnake), and *Notophthalmus viridescens* (Eastern Newt). Tadpole species included *Rana clamitans* (Green Frog) and *Lithobates catesbeianus* (Bullfrog).

### Experimental Design

The 14 ponds were divided equally into fertilized and unfertilized treatments. To increase nutrient concentrations in fertilized ponds, we added 0.1 L of 10-34-0 (N:P_2_O_5_) ammonium polyphosphate liquid fertilizer every two weeks from early July through September of 2021. This was equivalent to a loading rate of approximately 2.5 mg N/m^2^/d and 1.9 mg P/m^2^/d. This fertilization regime is based on recommendations for the management of small lakes and ponds from the Illinois DNR (Illinois Department of Natural Resources, 2017). The same regime was used in a different subset of ponds to show how nutrients influence the parasite load of largemouth bass (Stanley, et al., 2025). We measured the total water column nutrient ratios in ponds prior to assigning them to a treatment. Then, to minimize the likelihood of preexisting differences in pond nutrient ratios, we randomly assigned ponds to fertilized or control (unfertilized) treatments, until we achieved similar starting conditions across treatments.

### Sample Collection

We seined the length of each pond before treatment application and at the conclusion of the study to document fish species abundance and account for possible effects of fertilization on the trophic structure of the ponds. We used a six-foot depth seine with a 1/8-inch mesh. We recorded the number of individuals and sorted them into age classes (age 0-4+).

To assess the effects of nutrient additions on invertebrate communities and to account for any preexisting differences among pond communities, we collected invertebrates before treatment application and at the conclusion of the study. Initial sampling of invertebrates took place from mid-May to late-June. Ponds were then resampled for invertebrates from early-September to mid-October. During each sampling period, we interspersed our sampling of the ponds from each treatment temporally so that one haphazardly selected pond was sampled from each treatment before we sampled another replicate pond from the same treatment. To sample invertebrates, we did three rounds of benthic net sweeps at three depth points (0.5 m, 1 m, and the maximum wadeable depth) to collect surface-dwelling benthic invertebrates in each pond. Rounds lasted for one minute each, along a 2 m length of water. Because these ponds had loose muddy bottoms and we bobbed our net along the substrate in our sweeps, we suspect that we sampled a few inches into the substrate as well. For each pond, the three samples were combined and filtered through a 1 mm mesh sieve. All invertebrates were identified to family and sorted into FFGs using the classification from Merritt and colleagues (2005). Briefly, FFGs consist of five main categories that are used to infer resource use: scrapers that eat periphyton, shredders that eat CPOM, filtering-collectors (filterers) that eat FPOM, gathering-collectors (gatherers) that eat benthic FPOM, and predators that eat other animals. This classification provides information that allows generalization of the complex organization of aquatic communities (Mihuc 1997).

We measured dissolved oxygen (DO) and total C (TC), N (TN), and P (TP) every 6 or 7 days from June 28^th^ through August 17^th^. Samples for total water column nutrients were filtered through a small piece of acid-washed 60 μm mesh in the field. Ten dissolved oxygen loggers (MiniDOT, Precision Measurement Engineering) were rotated among the 14 ponds weekly so that dissolved oxygen and water temperature were recorded in each pond for ∼ 48 hours/week at 15-minute intervals. We attached the loggers to a buoy 1 meter below the surface.

We collected CPOM and FPOM after treatment application to clarify the link between our treatments and the diet of each FFG. To measure the suspended FPOM, we collected a 1L sample from each pond and filtered it onto glass fiber filters (pore size 0.7 um). We collected CPOM semi-quantitatively by completing two nets sweeps along the substrate over a 2 m length. Samples were filtered through a 1 mm sieve to separate CPOM from FPOM, and invertebrates were hand-picked out. Samples of CPOM and FPOM were oven-dried at 50° C until they achieved a stable weight for at least 24 hours.

### Nutrient Analyses

All particulate samples (CPOM, FPOM) were analyzed for particulate C and N using a Thermo Scientific Flash 2000 Elemental Analyzer in Southern Illinois University’s core facility for ecological analyses. Phosphorus was analyzed using the ascorbic acid molybdenum-blue method after digestion with potassium persulfate (APHA 1992). Weekly water column samples were measured on a Shimadzu total organic carbon/total nitrogen analyzer at Southern Illinois University’s core facility for ecological analyses.

### Metabolism estimates

Continuous measurements of dissolved oxygen concentrations and water temperature were collected from loggers to calculate pond metabolism. A weather station at Touch of Nature Environmental Center recorded wind speed, which was used to calculate gas exchange. Solar irradiance data, converted to photosynthetically active radiation (PAR), were collected from the Southern Illinois Airport weather station in Carbondale, IL.

Daily estimates of pond metabolism (GPP, R, and NEP) were calculated using the kalman model with the R package Lake Metabolizer (Winslow et al. 2016; LakeMetabolizer: *metab*.*kalman*). This metabolism model includes process error and fits parameters with maximum likelihood estimation using a Kalman filter (Kalman 1960, Harvey 1990). Estimation of metabolism using the kalman model is described in Winslow and colleagues (2016). The coefficient of gas exchange (*k*) was calculated using the Cole model (Winslow et al. 2016; LakeMetabolizer: *k*.*cole*) with wind speed data. The mixed layer depth (*z*) was estimated using water temperature data (LakeMetabolizer: *ts*.*meta*.*depths*), and saturated oxygen concentrations (O_s_) were calculated from temperature and altitude (LakeMetabolizer: *o2*.*at*.*sat*). We visually inspected the dissolved oxygen data for errors and removed all metabolism estimates where GPP was negative or R was positive.

### Statistical Analyses

We conducted all statistical analyses in R 4.5.2 (R Core Team 2025), and all ratios were natural log-transformed (ln-transformed) to avoid biases in interpretation (Isles 2020). We first quantified the effect of fertilization on pond nutrient concentrations and metabolism using generalized least squares regression. We regressed each nutrient and metabolism metric (TC, TN, TP, TC:TN, TC:TP, TN:TP, GPP, NEP, and R) against fertilization treatment nested within time (before or after fertilization began). These models included an autoregressive correlation structure (AR1) with time (week associated with a single day of year) nested within pond. One sample week (associated with the 195^th^ day of the year) between the pre- and post-treatment dates was excluded from this analysis to allow time for our treatment to take effect. We tested for fertilization effects on FPOM and CPOM concentration (g/L) and C:N, C:P, and N:P ratios by conducting Kruskal-Wallis rank sum tests on the difference between the pre and post estimates.

We used model selection to compare our ability to explain shifts in benthic community diversity, richness, the abundance of benthic invertebrates in each functional feeding group, zooplankton, and the abundance of two groups of zooplankton (cladocerans and copepods). We calculated Shannon diversity (Shannon 1948) and richness in each pond based on the number of individuals in each family using the package “vegan” in R (Dixon 2003). Shifts were calculated as the abundance at the end minus the abundance at the beginning of the experiment. These differences were rank transformed to avoid biases due to outliers or non-normal distributions. Because we were particularly interested in the effects of fertilization on the consumer assemblage, we restricted our candidate set to univariate models including fertilization and the mean estimate for each pond during the experiment for variables that responded to fertilization (Table 1). One pond was removed from all benthic invertebrate analyses, and one additional pond (two total) was removed from zooplankton analyses due to missing and unrecoverable data. We used Akaike’s Information Criterion (AICc) model selection corrected for small sample sizes and defined a top model as having a deltaAICc of <2 (Burnham and Anderson 2002). Finally, to assess the possibility that pre-existing communities of bluegill and green sunfish limited our ability to detect relationships between basal resources and invertebrates, we conducted model selection and analyzed top models once with the entire dataset, and then a second time with the (four) ponds with these fish populations excluded.

**Table 1:** Model coefficients and 95% confidence intervals indicating the estimated effect of time period (before or after fertilization) and fertilization treatment nested within each time period for water column nutrient availability: Total carbon (TC), nitrogen (TN), and phosphorus (TP) in mg/L and molar TC:TN, TC:TP, and TN:TP ratios, and whole pond metabolism: gross primary production (GPP), net ecosystem production (NEP), and respiration (R). We used generalized least squares regression with an autoregressive correlation structure with time nested within pond. Coefficients in bold indicate confidence intervals for the nested effect of fertilization that do not overlap with zero.

| Response | Term | Estimate | Lower CI | Upper CI |
| --- | --- | --- | --- | --- |
| TC (mg/L) | Before | 19.12 | 17.38 | 20.85 |
|  | After | 19.94 | 18.60 | 21.29 |
|  | Before/Fert | -0.50 | -2.96 | 1.95 |
|  | After/Fert | 0.79 | -1.11 | 2.69 |
| TN (mg/L) | Before | 0.77 | 0.66 | 0.88 |
|  | After | 0.74 | 0.65 | 0.82 |
|  | Before/Fert | 0.03 | -0.13 | 0.18 |
|  | **After/Fert** | **0.16** | **0.03** | **0.28** |
| TP (mg/L) | Before | 0.10 | 0.07 | 0.14 |
|  | After | 0.13 | 0.10 | 0.16 |
|  | Before/Fert | 0.02 | -0.03 | 0.08 |
|  | **After/Fert** | **0.10** | **0.06** | **0.14** |
| TC:TN (molar) | Before | 3.40 | 3.29 | 3.49 |
|  | After | 3.49 | 3.41 | 3.56 |
|  | Before/Fert | -0.03 | -0.17 | 0.11 |
|  | After/Fert | -0.09 | -0.20 | 0.02 |
| TC:TP (molar) | Before | 6.24 | 6.07 | 6.40 |
|  | After | 5.97 | 5.84 | 6.10 |
|  | Before/Fert | -0.16 | -0.39 | 0.08 |
|  | **After/Fert** | **-0.43** | **-0.62** | **-0.25** |
| TN:TP (molar) | Before | 2.93 | 2.78 | 3.09 |
|  | After | 2.60 | 2.48 | 2.71 |
|  | Before/Fert | -0.11 | -0.33 | 0.10 |
|  | **After/Fert** | **-0.31** | **-0.47** | **-0.14** |
| GPP (g O<sub>2</sub> m<sup>-2</sup> d<sup>-1</sup>) | Before | 5.17 | 3.47 | 6.87 |
| GPP (g O<sub>2</sub> m<sup>-2</sup> d<sup>-1</sup>) | After | 6.84 | 5.43 | 8.26 |
|  | Before/Fert | -0.40 | -2.80 | 2.00 |
|  | **After/Fert** | **-3.50** | **-5.58** | **-1.42** |
| NEP (g O<sub>2</sub> m<sup>-2</sup> d<sup>-1</sup>) | Before | -1.49 | -2.27 | -0.70 |
|  | After | -1.12 | -1.78 | -0.47 |
|  | Before/Fert | 0.28 | -0.83 | 1.39 |
|  | After/Fert | 0.29 | -0.67 | 1.26 |
| ER (g O<sub>2</sub> m<sup>-2</sup> d<sup>-1</sup>) | Before | -6.65 | -8.62 | -4.69 |
|  | After | -7.97 | -9.60 | -6.33 |
|  | Before/Fert | 0.68 | -2.10 | 3.46 |
|  | **After/Fert** | **3.80** | **1.39** | **6.20** |

## Results

We found that ponds that received the fertilization treatment had higher TN, TP and ER and lower TC:TP, TN:TP, and GPP (Table 1, Figure 1). After fertilization began, fertilized ponds were elevated by 0.16 mg TN/L, 0.10 mg TP/L. During this experimental period, in fertilized ponds, ER was elevated by 3.8 g 0_2_ m^-2^ d^-1^ and GPP was reduced by -3.5 g 0_2_ m^-2^ d^-1^ compared to unfertilized ponds. We did not detect effects of fertilization on FPOM or CPOM quantity or quality (C:N, C:P, and N:P, Appendix 1, Table 2).

**Figure 1.**
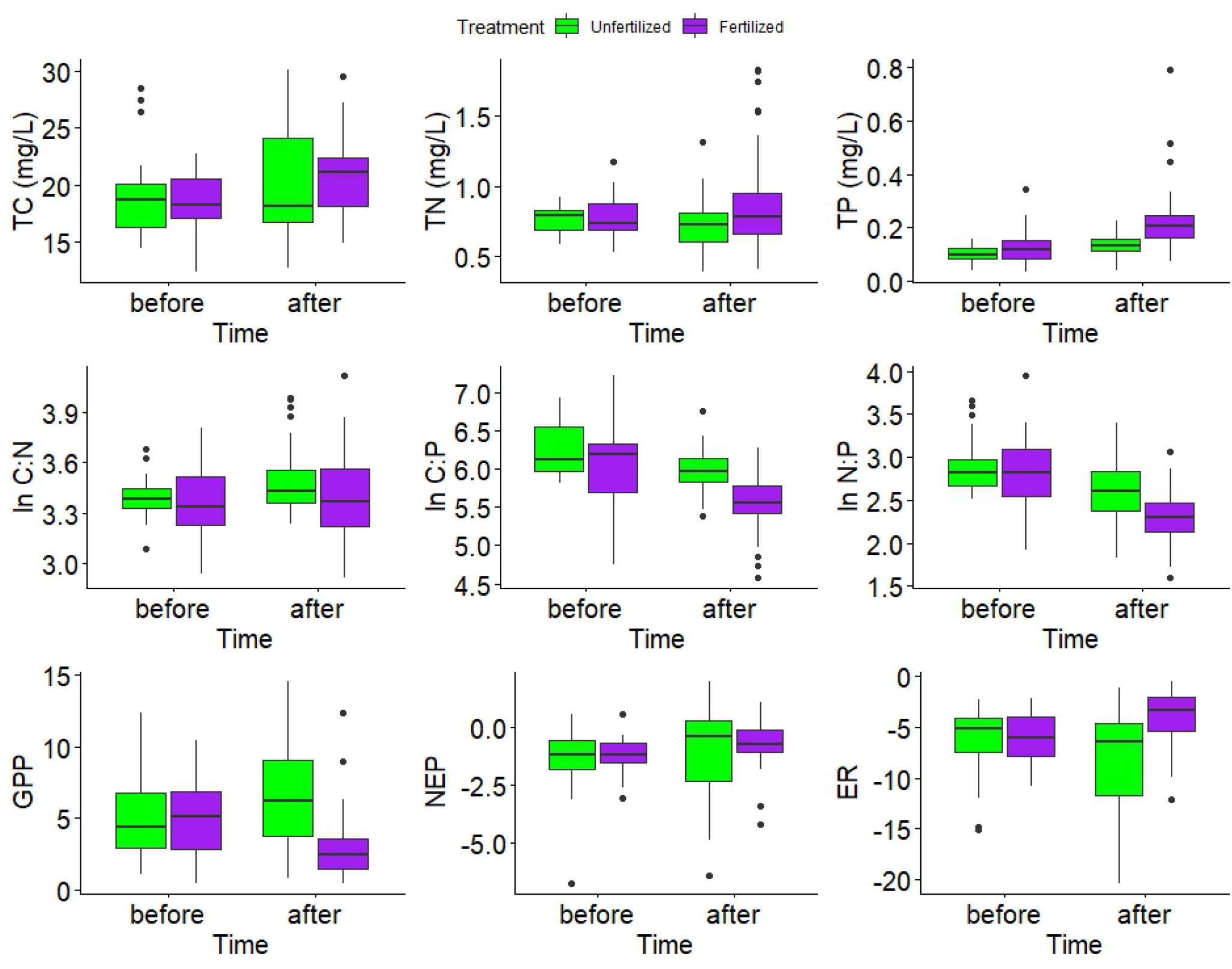
Total carbon (TC), nitrogen (TN), phosphorus (TP), natural log transformed molar ratios of these nutrient concentrations, and gross primary production (GPP), net ecosystem production (NEP), and ecosystem respiration (all estimated as g O_2_ m^-2^ d^-1^) in the water column of 7 fertilized (purple) and 7 unfertilized (green) ponds before and during (after) a fertilization experiment (Table 1). Water column nutrient ratios were manipulated by adding P-rich fertilizer every 2 weeks from early July through mid August.

Shifts in the Shannon index and richness of the benthic communities over time were best explained by the average natural log transformed TN:TP of the water column during the fertilization period (Appendix 1, Table 1). Greater increases in the Shannon indices and richness of ponds over the fertilization period were associated with lower TN:TP ratios (H: β = −13.85, t□ □ = −5.48, p < 0.001, R_2_ = 0.73; Richness: β =-10.02, t□ □=-2.70, p = 0.020, R^2^ = 0.40, Figure 2). Shifts in the abundance of cladocerans over time were also best explained by average pond TN:TP ratios during the experimental period (Appendix 1, Table 1); however, unlike benthic diversity, the abundance of cladocerans increased with TN:TP ratios (β = 10.55, t_10_ =2.85, p = 0.0173, R^2^=0.45, Figure 2). Shifts in the abundance of total benthic invertebrates and predators, in particular, over time were best explained by the average GPP and ER during the fertilization period (Appendix 1 Table 1). In addition, shifts in total benthic invertebrate abundances were equally well explained by a significant positive relationships with TN:TP ratios (β = 8.36, t□ □ =2.00, p = 0.071, R^2^=0.27) and a marginal negative effect of fertilization treatment (β = -3.71, t□ □=-1.89, p = 0.086, R^2^=0.24). Increased abundances of both benthic invertebrates and predators were associated with higher average GPP during the experimental period (benthic invertebrates: β = 0.64, t□ □ =2.42, p = 0.034, R^2^=0.35; predators: β = 0.82, t□ □ =3.87, p = 0.0026, R^2^=0.58, Figure 3) and lower average ER (benthic invertebrates: β = -.50, t□ □ =-2.14, p = 0.056, R^2^=0.29; predators: β =-0.66, t□ □ = -3.45, p = 0.0055, R^2^= 0.52, Figure 3). The null model was within delta AICc of 2 for shifts in all other FFGs (Appendix 1, Table 1). For all the analyses, models including fertilization alone were consistently ranked low (delta AIC > 2 for each regression).

**Figure 2.**
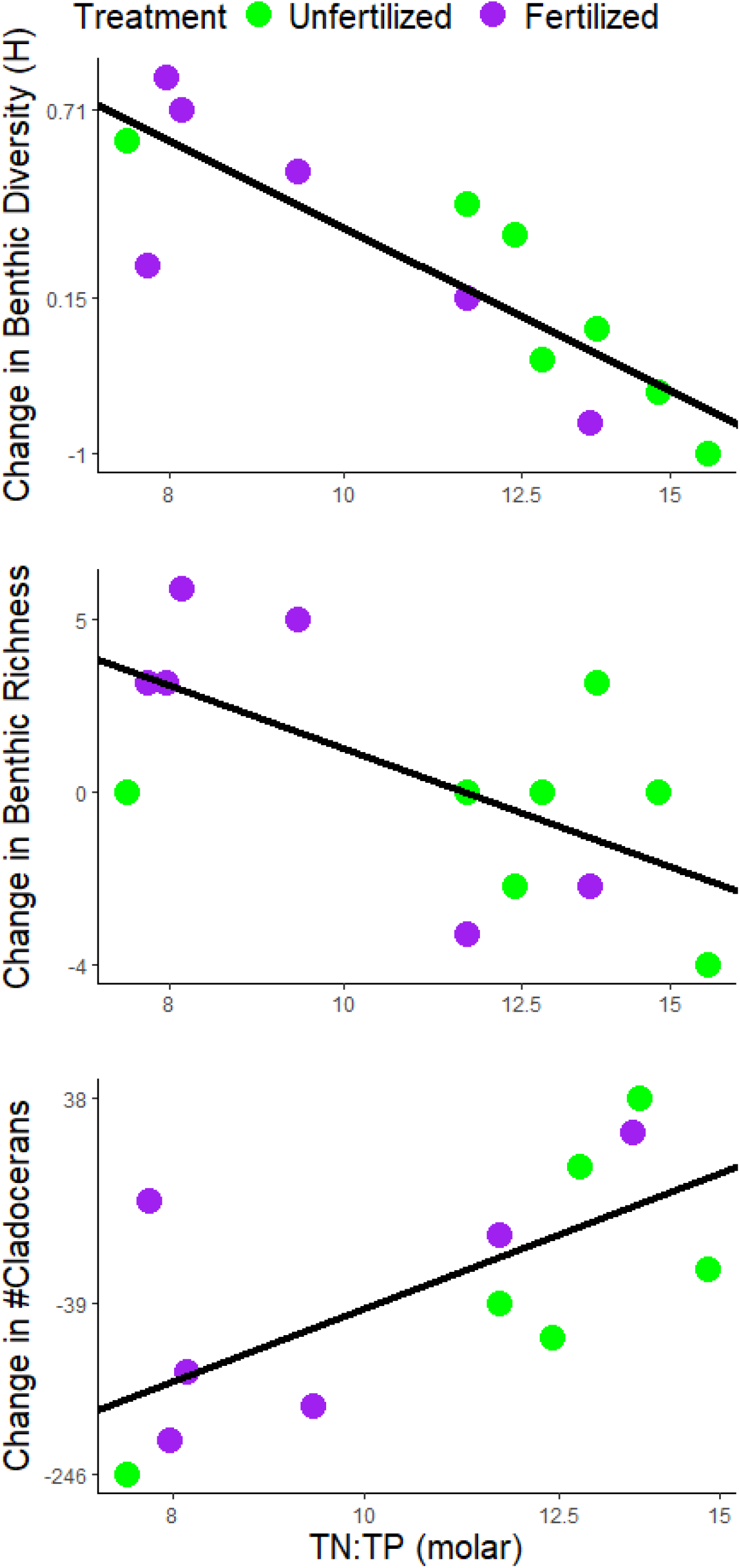
Change in benthic invertebrate Shannon index, family richness, and the abundance of cladocerans in 6 fertilized and 7 unfertilized experimental period as function of natural log transformed molar total nitrogen:phosphorus ratios (ln TN:TP) of the water column. Water column nutrient ratios were manipulated by adding P-rich fertilizer every 2 weeks from early July through mid August. The average ratios over this time period are shown. Note that the y axis has been rank transformed but displays that actual y values for each response.

**Figure 3.**
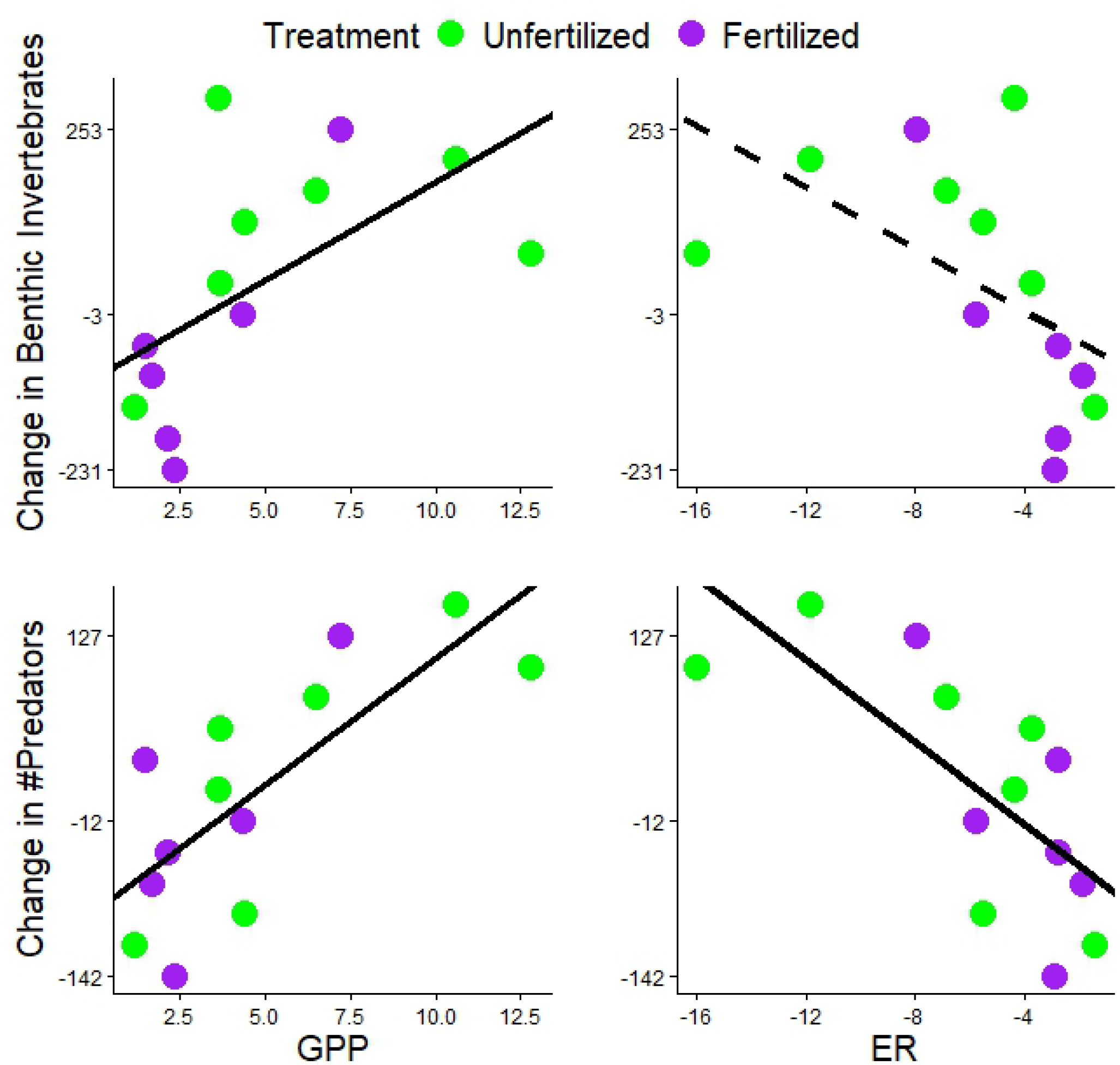
Change in the abundance of predators in 6 fertilized and 7 unfertilized experimental period as function of gross primary production (GPP, g O_2_ m^-2^ d^-1^) and ecosystem respiration (ER, g O_2_ m^-2^ d^-1^) of the water column. Water column nutrient ratios were manipulated by adding P-rich fertilizer every 2 weeks from early July through mid August. Note that the y axis has been rank transformed but displays that actual y values for each response.

Excluding ponds with pre-existing bluegill and green sunfish populations did not alter the direction of any of the above relationships, but it did cause the null model for shifts in benthic family richness and total benthic invertebrate abundance to be ranked within the top models (Appendix 1, Table 3). Excluding ponds with fish improved how well TN:TP ratios explained shifts in the Shannon index of benthic invertebrates from 73% to 86% (β = -11.78, t_7_ =-6.57, p < 0.001). However, in this subset of seven ponds, TP became the top model explaining shifts in diversity and explained 95% of this variation (β = 66.08, t_7_ =11.56, p < 0.001, Appendix 1, Table 3). Excluding ponds with fish also caused TN:TP ratios and TP concentrations in ponds to become the top model for shifts in the abundance of zooplankton (Appendix 1, Table 3). Increases in the abundance of zooplankton in this subset of ponds were positively associated with TN:TP ratios (β =9.89, t_7_ = 3.29, p = 0.013, R^2^= 0.61) and negatively associated with TP (β =-51.78, t_7_ = -3.13, p = 0.017, R^2^= 0.58). For all other response variables, excluding ponds with fish had no effect on the outcome of model selection. Excluding ponds with fish also improved how well GPP explained variation in shifts in predator abundances across ponds from 58%-68%, and the % of variation in shifts in cladoceran abundances explained by TN:TP increased from 45% to 72%.

## Discussion

Nutrient enrichment, primary production, and consumer assemblages are intricately linked through complex interactions (Hairston et al. 1960, Tilman 1982, Carpenter et al. 1985). Our experiment provides new evidence that consumer-mediated processes can influence the interaction between nutrient availability and production in semi-natural pond ecosystems. We found that fertilization reduced pond water column TN:TP ratios, GPP, and ER. These patterns indicate that fertilization likely caused or exacerbated N limitation in ponds, but this limitation did not appear to negatively affect ecosystem health: lower pond TN:TP ratios were associated with greater increases in benthic diversity over time. Additional lines of evidence suggest that, in addition to fertilization, shifts in the abundance of specific consumer groups may also have influenced TN:TP ratios, GPP, and ER. Consistent with predictions based on consumer-driven nutrient cycling, we found that ponds with lower TN:TP ratios were those that experienced bigger declines in the abundance of cladocerans over time. In addition, ponds with elevated GPP and ER during the experiment experienced bigger increases in the abundance of benthic predators over time, suggesting top-down control. Finally, each of these links between nutrient availability, GPP, ER, and consumers was stronger when ponds with pre-existing fish (bluegill and green sunfish) were removed from our analyses. This pattern underscores the importance of context and ecosystem complexity in mediating the role of invertebrate consumers in ecosystems.

The negative relationship between water column TN:TP ratios and both shifts in Shannon diversity and richness that we observed is consistent with positive effects of moderate nutrient enrichment observed in many other studies (Rader & Richardson 1992, Blumenshine et al. 1997, Miracle et al. 2006). While water column TN:TP ratios were a better predictor of Shannon diversity and richness than fertilization, fertilization reduced TN:TP ratios over time. Consistent with our observations, a mesocosm study found that intermediate fertilization increased invertebrate density and biomass (Miracle et al. 2006), and a study in the Everglades found that macroinvertebrate diversity and density were higher in enriched compared to unenriched areas (Rader & Richardson 1992). Likewise, a study of more than 250 lakes along a nutrient gradient in a eutrophying basin showed that both zooplankton and zoobenthos diversity peaked at intermediate nutrient levels (Wang et al. 2021). In general, it appears that invertebrate taxonomic richness follows a unimodal, bell-shaped pattern along a nutrient enrichment gradient, with richness decreased with heavy nutrient loading (Jeppesen et al. 2000, Wang et al. 2021). It’s important to note that several studies show declines in invertebrate diversity or density in response to heavy nutrient loading (Heino 2008, Yuan 2010, Schmera et al. 2017). A meta-analysis by Nessel and colleagues (2021) found that N and P additions typically decreased the diversity and abundance of invertebrates and that combined N and P enrichment produces stronger declines in aquatic ecosystems than N or P enrichment alone. Our study was conducted over a 2-month time frame; however, the effects of nutrient enrichment over a longer time frame may be different because of cumulative or long-term effects of nutrient enrichment.

While the relationship between water column TN:TP ratios that we observed indicates that our fertilization regime was not harmful (Knud-Hansen 1998, Illinois Department of Natural Resources 2017), we found that fertilization reduced rates of GPP and ER instead of increasing them (Figure 1). This is consistent with the possibility that many ponds were or became N-limited in our experiment. We observed increases in both the TN and TP of fertilized ponds, but the low N:P ratio of our fertilizer likely exacerbated N limitation. The average N:P ratios of the ponds during the two weeks before the experiment varied between 10 and 33 and, during the experiment, they declined to ratios ranging between 8.5 and 17. TN:TP ratios are imperfect predictors of nutrient limitation (Ptacnik et al. 2010), but these ratios are generally consistent with N limitation (Guildford and Hecky 2000).

In addition to fertilization, our results suggest that large declines in the abundance of cladocerans in some ponds also may have contributed to low water column TN:TP ratios. Compared to copepods (also in high abundance in our ponds), cladocerans have relatively low biomass N:P ratios (Walve and Larsson 1999), making their growth more sensitive to P limitation than N limitation. If low seston N:P ratios stimulated cladoceran population growth, we would expect a negative correlation between TN:TP and shifts in cladoceran abundance over time. However, we found that ponds with the lowest TN:TP ratios experienced the greatest declines in cladoceran abundance over time (Figure 2). Cladoceran abundance declined in most of the experimental ponds, but the extent of this decline varied dramatically among ponds. This is consistent with other work showing that zooplankton can act as P sinks over time (Urabe et al. 1995). Thus, we suspect that the loss of cladoceran biomass liberated phosphorus to the water column and that cladocerans moderated rather than unidirectionally responded to these shifts in nutrient availability.

Given that our nutrient additions likely had the unintended effect of exacerbating N limitation in these ponds, it may not be surprising that we found evidence supporting the control of GPP via benthic invertebrate predators, rather than support for clear bottom-up effects. Shifts in the abundance of invertebrate predators over time were best explained by pond GPP. GPP explained 58% of the variation in this shift across ponds and was positively associated with increased predator abundances. While our study does not test for these top down effects directly (e.g. through the exclusion of predators in some ponds), it is consistent with other studies showing that these top-down forces are apparent despite bottom-up variation in nutrient supply (Gilson and Mcquaid 2023). Trophic cascades can be triggered by a wide variety of predators including predacious benthic invertebrates (Malmqvist 1993, Atwood et al. 2013), but there is a bias toward the investigation of vertebrate predators, especially in aquatic ecosystems (Borer et al., 2005). Given the ubiquity of benthic invertebrate predators across ecosystems, and recent advancements in metabolism estimation (Appling et al. 2018, Jankowksi et al. 2021), we anticipate large gains in our understanding of ecosystem function from the continued quantification of the role of these less charismatic predators within energy flows.

The likelihood that the relationships we observed between predators and GPP (Figure 3) and caldocerans and nutrient ratios (Figure 2) were driven by variation in the abundance of these groups rather than bottom up processes is increased by our observation that these relationships were stronger when ponds with omnivorous fish populations were excluded from the analysis. The presence of omnivorous fish that feed on both primary producers and invertebrates is likely to obfuscate the relationship between basal resources (nutrient availability and production) and those of invertebrates whose diets have been carefully linked to their feeding morphology. Excluding these ponds improved how well GPP explained variation in shifts in predator abundances and how well TN:TP ratios explained shifts in cladoceran abundances \by 10 and 27 percentage points, respectively. Excluding these ponds also dramatically improved how well TN:TP ratios explained shifts in the Shannon index of benthic invertebrates (by 13 percentage points) and caused TP to explain 95% of the variation in this metric.

We did not observe any relationships between nutrients that responded to fertilization and benthic invertebrate functional feeding groups other than predators. In contrast with our results, N and P additions in streams decreased CPOM C:P ratios, leading to an increase in shredder biomass (Demi et al. 2019). Taken together, these contrasting results suggest that shredders might be more affected by changes in CPOM quality than quantity. Why our fertilization regime had no effect on CPOM quality is unclear, but we suspect that the CPOM in our ponds was comprised primarily of decomposing autochothonous macrophytes rather than the allochthonous material typically found in lotic ecosystems (these ponds were largely clear of tree cover).

The results of our experiment can inform our understanding of the role of small ponds, an important and often heavily managed ecosystem, within the carbon cycle. The ponds involved in this experiment were, on average, net heterotrophic (NEP < 0, Figure 1), indicating that all ponds were net sources of atmospheric carbon. Consistent with broad trends in lakes and estuaries (Hoellein et al. 2013), we found that, across all time points and ponds, gross primary production and ecosystem respiration were tightly correlated (adjusted R^2^ = 0.86, Appendix 1, Figure 1). As a result, fertilization and shifts in predator abundance were both associated with simultaneous shifts in rates of ER and GPP yielding no overall effect on net ecosystem production (Figures 1 and 3). The correlation between ER and GPP in our ponds suggests that respiration rates are largely driven by pelagic matter (Hoellein et al. 2013). Fertilization regimes designed to minimize pelagic respiration may improve our ability to store carbon in these heavily managed systems. We note that we specifically selected ponds with 3 trophic levels for our experiment. Many intentionally fertilized ponds are likely to have stocked piscivores serving as a 4^th^ trophic level. A concurrently run experiment using the same fertilization treatment in a different subset of experimental ponds with a 4^th^ trophic level (largemouth bass) showed that fertilization-induced declines in TN:TP ratios were linked to declines in the TN:TP ratios of a digestive tissue in largemouth bass which, in turn were linked to declines in the abundance of a trematode occupying this tissue (Stanley et al. 2025). Ultimately, quantifying links between nutrient availability, metabolic rates, and multiple endpoints of interest to land managers will facilitate our ability to enhance carbon sequestration in ponds.

## Acknowledgements

Thank you to all the field technicians that helped with this work: Morgan Brown, Anuj Pawar, Erin Sedlacek, Scott Binger, Sadie Edwards, Jonathan Henson, Tony Easton, and Caitlin Pelegrino. Amanda Rothert analyzed our carbon and nitrogen samples. Lusha Tronstad, Annika Walters, Jeff Baldock, Niall Clancy, Ashleigh Pilkerton, Margot Breiner, and Elizabeth Rieger provided comments that improved the manuscript.

## Notes

### Competing Interest Statement

The authors have declared no competing interest.

### Funding

- SIU Foundation Research Grant

## Literature Cited

Alimov, A. F. 2001. Studies on Biodiversity in the Plankton, Benthos, and Fish Communities, and the Ecosystems of Fresh Water Bodies Differing in Productivity 28:87–95.

APHA. 1992. Standard methods for the examination of water and wastewater. 18th edition. American Public Health Association, Washington, DC.

Appling, A. P., R. O. H. Jr, C. B. Yackulic, and M. Arroita. 2018. Overcoming EquifinalityL: Leveraging Long Time Series for Stream Metabolism Estimation. Journal of Geophysical Research: Biogeosciences 123:624–625.

Atkinson, C. L., K. A. Capps, A. T. Rugenski, and M. J. Vanni. 2017. Consumer-driven nutrient dynamics in freshwater ecosystemsL: from individuals to ecosystems 92:2003–2023.

Atwood, T. B., E. Hammill, H. S. Greig, P. Kratina, J. B. Shurin, D. S. Srivastava, and J. S. Richardson. 2013. Predator-induced reduction of freshwater carbon dioxide emissions. Nature Geoscience 6:191–194.

Balmer, M., and J. Downing. 2011. Carbon dioxide concentrations in eutrophic lakes: undersaturation implies atmospheric uptake. Inland Waters 1:125–132.

Brett, M. T., and C. R. Goldman. 1997. Consumer Versus Resource Control in Freshwater Pelagic Food Webs 275:16–19.

Burnham, K. P., and D. R. Anderson. 2002. Model Selection and Multimodel Inference: A Practical Information-Theoretic Approach. Springer New York, NY.

Carpenter, S. R., J. F. Kitchell, S. R. Carpenter, and J. F. Kitchell. 1988. Consumer Control of Lake Productivity 38:764–769.

Carpenter, S. R., J. F. Kitchell, and J. R. Hodgson. 1985. Cascading trophic interactions and lake productivity: fish predation and herbivory can regulate lake ecosystems. The American Institute of Biological Sciences 35:634–639.

Cummins, K. W., R. W. Merritt, and P. C. N. Andrade. 2005. The use of invertebrate functional groups to characterize ecosystem attributes in selected streams and rivers in south Brazil. Studies on Neotropical Fauna and Environment 40:69–89.

Dixon, P. 2003. VEGAN, a package of R functions for community ecology. Journal of Vegetation Science 14:927–930.

Downing, J. A., J. J. Cole, and W. H. Mcdowell. 2006. The global abundance and size distribution of lakes, ponds, and impoundments 51:2388–2397.

Elser, J. J., M. E. S. Bracken, E. E. Cleland, D. S. Gruner, W. S. Harpole, J. T. Ngai, E. W. Seabloom, J. B. Shurin, and J. E. Smith. 2007. Global analysis of nitrogen and phosphorus limitation of primary producers in freshwater, marine and terrestrial ecosystems. Ecology Letters 10:1–8.

Elser, J. J., and J. Urabe. 1999. The stoichiometry of consumer-driven nutrient recycling: Theory, observations, and consequences. Ecology 80:735–751.

Gilson, A. R., and C. Mcquaid. 2023. Top-down versus bottom-upL: Grazing and upwelling regime alter patterns of primary productivity in a warm-temperate system:1–17.

del Giorgio, P. A., and R. H. Peters. 1994. Patterns in Planktonic PL: R Ratios in LakesL: Influence of Lake Trophy and Dissolved Organic Carbon. Limnology and Oceanography 39:772–787.

Guildford, S. J., and R. E. Hecky. 2000. Total nitrogen, total phosphorus, and nutrient limitation in lakes and oceansL: Is there a common relationshipL? 45:1213–1223.

Hairston, N. G., F. E. Smith, and L. B. Slobodkin. 1960. Community Structure, Population Control, and Competition. The American Naturalist 94:421–425.

Halpern, B.S., S. Walbridge, K.A. Selkoe, C.V. Kappel, F. Micheli, C. d’Agrosa, J.F. Bruno, K.S. Casey, C. Ebert, H.E. Fox, and R. Fujita. 2008. A global map of human impact on marine ecosystems. Science, 319(5865): 948–952.

Hanson, P. C., D. L. Bade, S. R. Carpenter, T. K. Kratz, and W. Long. 2003. Lake metabolismL: Relationships with dissolved organic carbon and phosphorus 48:1112–1119.

Harvey, A.C. 1990. Forecasting, structural time series models and the Kalman filter. Camrbidge University Press.

Hecky, R. E., and P. Kilham. 1988. Nutrient limitation of phytoplankton in freshwater and marine environmentsL: A review of recent evidence on the effects of enrichment1 Nutritional requirements of phytoplankton Algal cells require elements in relatively 33:796–822.

Heino, J. 2008. Patterns of functional biodiversity and functionLenvironment relationships in lake littoral macroinvertebrates. Limnology and Oceanography. 53(4): 1446–1455.

Hoellein, T. J., D. A. Bruesewitz, D. C. Richardson, T. J. Hoellein, D. A. Bruesewitz, and D. C. Richardson. 2013. Revisiting Odum (1956): A synthesis of aquatic ecosystem metabolism. Limnologica 58:2089–2100.

Illinois Department of Natural Resources. Revised 2017. Management of Small Lakes and Ponds in Illinois. Third Edition.

Isles, P. D. F. 2020. The misuse of ratios in ecological stoichiometry. Ecology 101:1–7.

Jankowksi, K. J., F. H. Mejia, J. R. Blaszczak, and G. W. Holtgrieve. 2021. Aquatic ecosystem metabolism as a tool in environmental management. WIREs Water 8:e1521.

Jeppesen, E., Peder Jensen, J., SØndergaard, M., Lauridsen, T. and Landkildehus, F. 2000. Trophic structure, species richness and biodiversity in Danish lakes: changes along a phosphorus gradient. Freshwater biology, 45(2): 201–218.

Kalman, R.E. 1960. A new approach to linear filtering and prediction problems. Journal of Basic Engineering. 82:35–45.

Knud-Hansen, C.F, C. Fulton, and D. Clair. 1998. Pond fertilization: ecological approach and practical application. Corvallis, Oregon: Pond Dynamics/Aquaculture Collaborative Research Support Program, Oregon State University.

Malmqvist, B. 1993. Interactions in Stream Leaf PacksL: Effects of a Stonefly Predator on Detritivores and Organic Matter Processing. Oikos 66:454–462.

Matich, P., M. R. Heithaus, and C. A. Layman. 2011. Contrasting patterns of individual specialization and trophic coupling in two marine apex predators: 294–305.

Mihuc, T.B. 1997. The functional trophic role of lotic primary consumers: generalist versus specialist strategies. Freshwater biology, 37(2): 455–462.

Miracle, M.R., Moss, B., Vicente, E., Romo Pérez, S., Rueda, J., Bécares Mantecón, E., Fernández-Aláez, M.D.C., Fernández-Aláez, M., Hietala, J., Kairesalo, T. and Vakkilainen, K. 2006. Response of macroinvertebrates to experimental nutrient and fish additions in European localities at different latitudes. Limnetica. 25(1-2): 585–612.

Nessel, M.P., Konnovitch, T., Romero, G.Q. and González, A.L. 2021. Nitrogen and phosphorus enrichment cause declines in invertebrate populations: a global metaLanalysis. Biological Reviews. 96(6): 2617–2637.

Peñuelas, J., J. Sardans, A. Rivas-ubach, and I. A. Janssens. 2012. The human-induced imbalance between C, N and P in Earth’s life system. Global Change Biology 18:3–6.

Ptacnik, R., T. Andersen, and T. Tamminen. 2010. Performance of the Redfield Ratio and a Family of Nutrient Limitation Indicators as Thresholds for Phytoplankton N vs. P Limitation: 1201–1214.

Rader, R.B. and Richardson, C.J. 1992. The effects of nutrient enrichment on algae and macroinvertebrates in the Everglades: a review. Wetlands, 12(2): 121–135.

Rizzuto, M., S. J. Leroux, and O. J. Schmitz. 2024. Rewiring the Carbon CycleL: A Theoretical Framework for Animal L Driven Ecosystem Carbon Sequestration: 1–20.

Schindler, D. W., and E. J. Fee. 1974. Experimental Lakes Area: Whole-Lake Experiments in Eutrophication. Journal of the Fisheries Board of Canada 31:937–953.

Schmera, D., J. Heino, J. Podani, T. Eros, and S. Dolédec. 2017. Functional diversity: a review of methodology and current knowledge in freshwater macroinvertebrate research. Hydrobiologia 787:27–44.

Shannon, C.E. 1948. A mathematical theory of communication. The Bell system technical journal, 27(3): 379–423.

Staehr, P. A., D. Bade, M. C. van de Bogert, G. R. Koch, C. Williamson, P. Hanson, J. J. Cole, and T. Kratz. 2010. Lake metabolism and the diel oxygen technique: State of the science. Limnology and Oceanography: Methods 8:628–644.

Stanley, A., I. Sadler, S. Binger, and C. Narr. 2025. Host tissue stoichiometry links nutrient availability to parasite abundance in a piscivorous fish. Research Square (Preprint) 10.21203/rs.3.rs-7329746/v1

Tilman, D. 1982. Resource competition and community structure. Princeton University Press.

Urabe, J., M. Nakanishi, and K. Kawabata. 1995. Contribution of Metazoan Plankton to the Cycling of Nitrogen and Phosphorus in Lake Biwa. Limnology and Oceanography 40:232–241.

Vitousek, P. M., J. D. Aber, R. W. Howarth, G. E. Likens, A. Pamela, D. W. Schindler, W. H. Schlesinger, and D. G. Tilman. 1997. Human alteration of the global nutrogen cycle: sources and consequences. Ecological Applications 7:737–750.

Wang, H., Molinos, J.G., Heino, J., Zhang, H., Zhang, P. and Xu, J. 2021. Eutrophication causes invertebrate biodiversity loss and decreases cross-taxon congruence across anthropogenically-disturbed lakes. Environment International. 153:106494.

Walve, J., and U. Larsson. 1999. Carbon, nitrogen and phosphorus stoichiometry of crustacean zooplankton in the Baltic SeaL: implications for nutrient recycling 21:2309–2321.

Welti, N., M. Striebel, A. J. Ulseth, W. F. Cross, S. DeVilbiss, P. M. Glibert, L. Guo, A. G. Hirst, J. Hood, J. S. Kominoski, K. L. MacNeill, A. S. Mehring, J. R. Welter, and H. Hillebrand. 2017. Bridging food webs, ecosystem metabolism, and biogeochemistry using ecological stoichiometry theory. Frontiers in Microbiology 8:1–14.

Winslow, L. A., J. A. Zwart, R. D. Batt, H. A. Dugan, R. Iestyn, J. R. Corman, P. C. Hanson, J.S. Read, L. A. Winslow, J. A. Zwart, R. D. Batt, H. A. Dugan, R. Iestyn, J. R. Corman, P.C. Hanson, J. S. Read, L. A. Winslow, J. A. Zwart, R. D. Batt, H. A. Dugan, R. I. Woolway, R. Jessica, P. C. Hanson, and J. S. Read. 2016. LakeMetabolizerL: an R package for estimating lake metabolism from free-water oxygen using diverse statistical models LakeMetabolizerL: an R package for estimating lake metabolism from free-water oxygen using diverse statistical models. Inland Waters 6:622–636.

Yuan, L.L. 2010. Estimating the effects of excess nutrients on stream invertebrates from observational data. Ecological Applications. 20(1): 110–125.

