## Appendix 1 for "Shifts in consumer assemblages are linked to nutrient availability and ecosystem metabolism"

Isabella G. Sadler, Adrienne L. Stanley, and Charlotte F. Narr **Shifts in consumer assemblages are linked to nutrient availability and ecosystem metabolism**

Table 1: Model selection for linear models predicting shifts in benthic invertebrate community diversity (Shannon 1948), richness, the abundance of benthic invertebrates in multiple functional feeding groups (Cummins et al., 2025), and the abundance of two groups of zooplankton (cladocerans and copepods) over an approximately 3 month long fertilization period. Shifts were calculated as the abundance at the end minus the abundance at the beginning of the fertilization period. These differences were rank transformed to avoid biases due to outliers or non-normal distributions. Number of parameters (K), change in AIC_C_ compared to the best-ranked model (ΔAIC_C_), Akaike model weights (*W*), and log likelihood estimate (*LL*) for top models (ΔAIC_C_ < 2) are shown in bold.

| **Response** | **Model Predictor** | **K** | **ΔAIC_C_** | ***W*** | ***LL*** |
| --- | --- | --- | --- | --- | --- |
| **Benthic Invertebrates** | **GPP** | **3** | **0.00** | **0.29** | **-32.82** |
|  | **ER** | **3** | **1.04** | **0.18** | **-33.34** |
|  | **ln TN:TP** | **3** | **1.53** | **0.14** | **-33.59** |
|  | **Fert** | **3** | **1.91** | **0.11** | **-33.77** |
|  | Null | 2 | 2.09 | 0.10 | -35.60 |
|  | ln TC:TP | 3 | 2.18 | 0.10 | -33.91 |
|  | TP | 3 | 3.25 | 0.06 | -34.44 |
|  | TN | 3 | 5.56 | 0.02 | -35.60 |
| **Cladocerans** | **ln TN:TP** | **3** | **0.00** | **0.65** | **-28.33** |
|  | Null | 2 | 3.46 | 0.12 | -31.89 |
|  | ln TC:TP | 3 | 3.55 | 0.11 | -30.11 |
|  | TP | 3 | 6.13 | 0.03 | -31.39 |
|  | TN | 3 | 6.29 | 0.03 | -31.48 |
|  | ER | 3 | 6.96 | 0.02 | -31.81 |
|  | GPP | 3 | 7.00 | 0.02 | -31.83 |
|  | Fert | 3 | 7.01 | 0.02 | -31.84 |
| Copepods | TP | 3 | 0.00 | 0.24 | -30.01 |
|  | Null | 2 | 0.11 | 0.22 | -31.89 |
|  | lnTC:TP | 3 | 0.35 | 0.20 | -30.18 |
|  | TN | 3 | 1.48 | 0.11 | -30.75 |
|  | ER | 3 | 2.50 | 0.07 | -31.26 |
|  | GPP | 3 | 2.86 | 0.06 | -31.44 |
|  | Fert | 3 | 3.05 | 0.05 | -31.53 |
|  | TN:TP | 3 | 3.17 | 0.05 | -31.59 |
| **Diversity (H)** | **ln TN:TP** | **3** | **0.00** | **0.99** | **-27.04** |
|  | ln TC:TP | 3 | 12.37 | 0.00 | -33.23 |
|  | Null | 2 | 13.65 | 0.00 | -35.60 |
|  | TP | 3 | 15.25 | 0.00 | -34.67 |
|  | TN | 3 | 15.36 | 0.00 | -34.72 |
|  | Fert | 3 | 15.62 | 0.00 | -34.85 |
|  | GPP | 3 | 16.12 | 0.00 | -35.10 |
|  | ER | 3 | 16.22 | 0.00 | -35.15 |
| Filtering  Collectors | ln TN:TP | 3 | 0.00 | 0.41 | -32.40 |
|  | Null | 2 | 1.54 | 0.19 | -34.90 |
|  | TN | 3 | 3.27 | 0.08 | -34.04 |
|  | ER | 3 | 3.31 | 0.08 | -34.06 |
|  | Fert | 3 | 3.33 | 0.08 | -34.06 |
|  | GPP | 3 | 3.33 | 0.08 | -34.06 |
|  | ln TC:TP | 3 | 3.89 | 0.06 | -34.35 |
|  | TP | 3 | 4.90 | 0.04 | -34.85 |
| Gathering Collectors | Null | 2 | 0.00 | 0.41 | -35.60 |
|  | ln TC:TP | 3 | 2.71 | 0.11 | -35.22 |
|  | Fert | 3 | 2.90 | 0.10 | -35.32 |
|  | ln TN:TP | 3 | 2.99 | 0.09 | -35.36 |
|  | TP | 3 | 3.35 | 0.08 | -35.54 |
|  | TN | 3 | 3.44 | 0.07 | -35.59 |
|  | GPP | 3 | 3.45 | 0.07 | -35.59 |
|  | ER | 3 | 3.47 | 0.07 | -35.60 |
| Zooplankton | Null | 2 | 0.00 | 0.38 | -31.89 |
|  | ln TC:TP | 3 | 2.48 | 0.11 | -31.30 |
|  | TP | 3 | 2.60 | 0.10 | -31.36 |
|  | ln TN:TP | 3 | 2.80 | 0.09 | -31.46 |
|  | ER | 3 | 2.83 | 0.09 | -31.48 |
|  | GPP | 3 | 2.98 | 0.08 | -31.55 |
|  | TN | 3 | 3.28 | 0.07 | -31.70 |
|  | Fert | 3 | 3.41 | 0.07 | -31.77 |
| **Predators** | **GPP** | **3** | **0.00** | **0.66** | **-30.01** |
|  | **ER** | **3** | **1.66** | **0.29** | **-30.84** |
|  | ln TN:TP | 3 | 7.25 | 0.02 | -33.64 |
|  | Null | 2 | 7.71 | 0.01 | -35.60 |
|  | TP | 3 | 8.75 | 0.01 | -34.39 |
|  | ln TC:TP | 3 | 9.80 | 0.00 | -34.91 |
|  | Fert | 3 | 10.35 | 0.00 | -35.19 |
|  | TN | 3 | 11.16 | 0.00 | -35.59 |
| **Richness** | **ln TN:TP** | **3** | **0.00** | **0.61** | **-32.01** |
|  | Null | 2 | 3.16 | 0.12 | -35.33 |
|  | ln TC:TP | 3 | 3.90 | 0.09 | -33.96 |
|  | Fert | 3 | 4.85 | 0.05 | -34.44 |
|  | TP | 3 | 5.57 | 0.04 | -34.80 |
|  | TN | 3 | 5.58 | 0.04 | -34.80 |
|  | ER | 3 | 6.17 | 0.03 | -35.10 |
|  | GPP | 3 | 6.31 | 0.03 | -35.17 |
| Scrapers | GPP | 3 | 0.00 | 0.37 | -32.90 |
|  | ER | 3 | 0.24 | 0.33 | -33.02 |
|  | Null | 2 | 1.90 | 0.14 | -35.58 |
|  | Fert | 3 | 4.06 | 0.05 | -34.93 |
|  | ln TC:TP | 3 | 4.77 | 0.03 | -35.28 |
|  | ln TN:TP | 3 | 5.17 | 0.03 | -35.48 |
|  | TN | 3 | 5.17 | 0.03 | -35.48 |
|  | TP | 3 | 5.36 | 0.03 | -35.58 |

Table 2: Results of Kruskal-Wallis tests comparing the change in CPOM and FPOM molar nutrient ratios (carbon(C):nitrogen (N), C:phosphorus (P), and N:P) and density over a 3 month fertilization experiment in 6 fertilized and 7 unfertilized experimental ponds.

|  | **Response** | **Chi-squared** | **p-value** |
| --- | --- | --- | --- |
| CPOM | C:P | 0.0066 | 0.94 |
|  | C:N | 0.33 | 0.57 |
|  | N:P | 0.16 | 0.68 |
|  | Density (g/L) | 0.73 | 0.39 |
| FPOM | C:P | 0.33 | 0.57 |
|  | C:N | 2.47 | 0.12 |
|  | N:P | 1 | 0.32 |
|  | Density (g/L) | 0.18 | 0.67 |

Table 3: Model selection for linear models excluding ponds where fish were present predicting shifts in benthic invertebrate community diversity (Shannon 1948), richness, the abundance of benthic invertebrates in multiple functional feeding groups (Cummins et al., 2025), and the abundance of two groups of zooplankton (cladocerans and copepods) over an approximately 3 month long fertilization period. Shifts were calculated as the abundance at the end minus the abundance at the beginning of the fertilization period. These differences were rank transformed to avoid biases due to outliers or non-normal distributions. Number of parameters (K), change in AIC_C_ compared to the best-ranked model (ΔAIC_C_), Akaike model weights (*W*), and log likelihood estimate (*LL*) for top models (ΔAIC_C_ < 2) are shown in bold. Response variables were modeled based on average water column measurements during the fertilization period that responded to fertilization. These included total total nitrogen (TN) total phosphorus (TP). natural log transformed molar ratios of total carbon:TP (ln TC:TP) and TN:TP, gross primary production (GPP, g O_2_ m^-2^ d^-1^), and ecosystem respiration (ER, g O_2_ m^-2^ d^-1^).

| **Response** | **Model Predictor** | **K** | **ΔAIC_C_** | ***W*** | ***LL*** |
| --- | --- | --- | --- | --- | --- |
| Benthic Invertebrates | Null | 2 | 0.00 | 0.44 | -21.31 |
|  | ln TN:TP | 3 | 2.79 | 0.11 | -20.30 |
|  | ln TC:TP | 3 | 3.11 | 0.09 | -20.46 |
|  | TN | 3 | 3.25 | 0.09 | -20.53 |
|  | GPP | 3 | 3.43 | 0.08 | -20.62 |
|  | Fert | 3 | 3.51 | 0.08 | -20.66 |
|  | ER | 3 | 3.83 | 0.06 | -20.82 |
|  | TP | 3 | 4.35 | 0.05 | -21.08 |
| **Cladocerans** | **ln TN:TP** | **3** | **0.00** | **0.54** | **-15.58** |
|  | **TP** | **3** | **1.05** | **0.32** | **-16.10** |
|  | ln TC:TP | 3 | 3.22 | 0.11 | -17.19 |
|  | Null | 2 | 6.66 | 0.02 | -21.31 |
|  | Fert | 3 | 10.76 | 0.00 | -20.96 |
|  | TN | 3 | 10.83 | 0.00 | -20.99 |
|  | GPP | 3 | 11.19 | 0.00 | -21.17 |
|  | ER | 3 | 11.27 | 0.00 | -21.21 |
| Copepod | ln TN:TP | 3 | 0.00 | 0.34 | -18.03 |
|  | TP | 3 | 0.24 | 0.30 | -18.15 |
|  | Null | 2 | 1.75 | 0.14 | -21.31 |
|  | ln TC:TP | 3 | 2.23 | 0.11 | -19.15 |
|  | ER | 3 | 4.19 | 0.04 | -20.13 |
|  | GPP | 3 | 4.69 | 0.03 | -20.38 |
|  | Fert | 3 | 5.26 | 0.02 | -20.66 |
|  | TN | 3 | 6.26 | 0.01 | -21.17 |
| **Diversity (H)** | **TP** | **3** | **0.00** | **0.99** | **-7.81** |
|  | ln TN:TP | 3 | 9.26 | 0.01 | -12.44 |
|  | ln TC:TP | 3 | 19.35 | 0.00 | -17.49 |
|  | Null | 2 | 22.20 | 0.00 | -21.31 |
|  | TN | 3 | 23.45 | 0.00 | -19.53 |
|  | Fert | 3 | 24.89 | 0.00 | -20.26 |
|  | GPP | 3 | 24.93 | 0.00 | -20.27 |
|  | ER | 3 | 25.18 | 0.00 | -20.40 |
| Filtering collectors | Null | 2 | 0.00 | 0.34 | -20.92 |
|  | Fert | 3 | 1.91 | 0.13 | -19.47 |
|  | ER | 3 | 2.28 | 0.11 | -19.66 |
|  | GPP | 3 | 2.37 | 0.10 | -19.70 |
|  | TP | 3 | 2.49 | 0.10 | -19.76 |
|  | ln TN:TP | 3 | 2.51 | 0.10 | -19.77 |
|  | ln TC:TP | 3 | 3.13 | 0.07 | -20.08 |
|  | TN | 3 | 3.87 | 0.05 | -20.45 |
| Gathering collectors | Null | 2 | 0.00 | 0.55 | -21.31 |
|  | ln TC:TP | 3 | 3.56 | 0.09 | -20.69 |
|  | Fert | 3 | 4.10 | 0.07 | -20.96 |
|  | TN | 3 | 4.31 | 0.06 | -21.06 |
|  | ER | 3 | 4.40 | 0.06 | -21.11 |
|  | ln TN:TP | 3 | 4.42 | 0.06 | -21.12 |
|  | GPP | 3 | 4.56 | 0.06 | -21.19 |
|  | TP | 3 | 4.71 | 0.05 | -21.26 |
| **Predator** | **GPP** | **3** | **0.00** | **0.66** | **-16.15** |
|  | ER | 3 | 2.04 | 0.24 | -17.17 |
|  | Null | 2 | 5.52 | 0.04 | -21.31 |
|  | ln TN:TP | 3 | 5.90 | 0.03 | -19.10 |
|  | TP | 3 | 7.99 | 0.01 | -20.14 |
|  | ln TC:TP | 3 | 9.77 | 0.00 | -21.03 |
|  | Fert | 3 | 10.01 | 0.00 | -21.15 |
|  | TN | 3 | 10.31 | 0.00 | -21.30 |
| Richness | Null | 2 | 0.00 | 0.43 | -20.87 |
|  | TN | 3 | 1.98 | 0.16 | -19.47 |
|  | TP | 3 | 2.54 | 0.12 | -19.74 |
|  | ln TC:TP | 3 | 3.08 | 0.09 | -20.02 |
|  | ln TN:TP | 3 | 3.88 | 0.06 | -20.42 |
|  | Fert | 3 | 4.27 | 0.05 | -20.61 |
|  | ER | 3 | 4.60 | 0.04 | -20.78 |
|  | GPP | 3 | 4.76 | 0.04 | -20.85 |
| Scraper | Null | 2 | 0.00 | 0.39 | -21.31 |
|  | GPP | 3 | 1.44 | 0.19 | -19.63 |
|  | ER | 3 | 1.76 | 0.16 | -19.79 |
|  | ln TC:TP | 3 | 2.95 | 0.09 | -20.38 |
|  | TN | 3 | 4.04 | 0.05 | -20.93 |
|  | Fert | 3 | 4.10 | 0.05 | -20.96 |
|  | TP | 3 | 4.58 | 0.04 | -21.20 |
|  | ln TN:TP | 3 | 4.76 | 0.04 | -21.29 |
| **Zooplankton** | **ln TN:TP** | **3** | **0.00** | **0.45** | **-17.11** |
|  | **TP** | **3** | **0.52** | **0.35** | **-17.37** |
|  | Null | 2 | 3.60 | 0.07 | -21.31 |
|  | ln TC:TP | 3 | 4.68 | 0.04 | -19.45 |
|  | ER | 3 | 4.93 | 0.04 | -19.58 |
|  | GPP | 3 | 5.35 | 0.03 | -19.78 |
|  | Fert | 3 | 7.11 | 0.01 | -20.66 |
|  | TN | 3 | 7.93 | 0.01 | -21.08 |

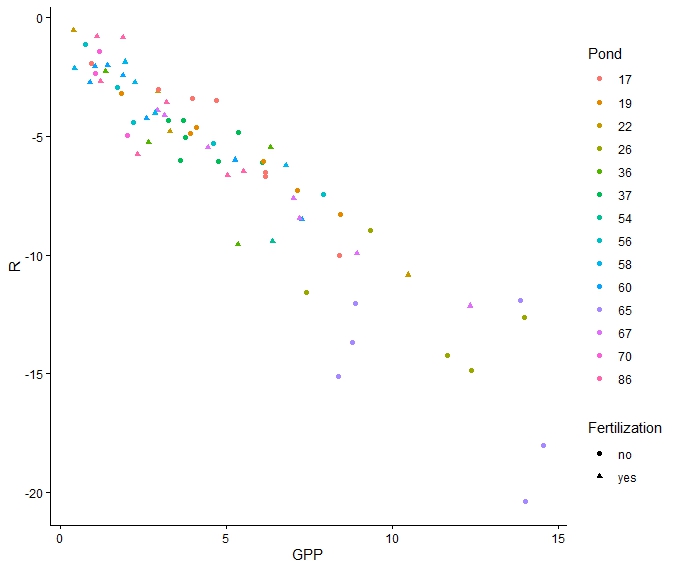

Figure 1: Relationship between ecosystem respiration rates (R) and gross primary production (GPP) in 14 experimental ponds before and during a 2 month fertilization experiment. Ponds are shown differentiated in different colors and fertilization treatment is shown in shapes. Generalized least squares regression of GPP against R indicates a significant relationship (-0.81, t=-21.12, p < 0.001, Adjusted R^2^ = 0.86). This model included an autoregressive correlation structure (AR1) with time (week associated with a single day of year) nested within pond.
